# Molecular mechanisms of dynamics-mediated effects of pathogenic missense mutations on Menin protein function

**DOI:** 10.1101/2024.12.22.630010

**Authors:** Qian Zhang, Hao Wang, Xiaohui Chen, Wenjian Li, Jin Peng, Manjie Zhang, Bin Sun

## Abstract

Understanding how disease-causing missense mutations (DCMMs) affect protein function is fundamental. The traditional structure-function paradigm for proteins has evolved into a structure-dynamics-function model, as dynamics is increasingly recognized as a key regulator of protein function. Consequently, it is essential to incorporate dynamics into the once heavily emphasized structure-function framework to explain the effects of DCMMs. Although research in this area is emerging, evidence supporting a definitive role of dynamics in mediating DCMM effects on protein function remains limited. In this study, we used Menin—a mutation-prone scaffold protein involved in various pathologies—as a model system to explore how DCMMs affect Menin’s function. By performing molecular dynamics (MD) simulations on 24 clinically confirmed DCMMs, we showed that DCMMs do not necessarily destabilize protein stability. Instead, they induce similar dynamic changes in the protein. The consequences of DCMMs on Menin’s function were further assessed through umbrella sampling of the Menin-JunD protein protein interaction (PPI). We found that DCMMs attenuate this interaction and disrupt a highly conserved JunD dissociation pathway in wild-type (WT) Menin. The underlying mechanism was revealed through allosteric analysis, which showed that, despite being located far from the JunD binding site, DCMMs uniformly disturbed the coupling between residue E179 and the binding pocket. Additionally, forced maintenance of E197-pocket coupling restored the impaired Menin-JunD PPI in DCMMs. Together, these data demonstrate that DCMMs alter Menin’s function through dynamics, with allostery playing a crucial role. This study further supports the incorporation of dynamics into the traditional ”structure-function” diagram to better explain the effects of DCMMs.

## 1 Introduction

Understanding the molecular mechanisms by which disease-causing missense mutations (DCMM) alter protein function is fundamental to the development of therapeutic strategies. In the early stages, the availability of protein structures from the PDB databank sparked investigations into the effects of mutations on structural stability, often based on analyses of structural quality—such as hydrogen bonding and side-chain packing—or on thermostability analyses, like calculating protein folding free energy [1, 2]. Emphasis was later placed on the conformational stability of proteins to explain the effects of disease-causing mutations, which fostered the notion that these mutations are more likely to destabilize protein conformation [2, 3]. However, relying solely on conformational stability cannot explain the effects of all disease-causing mutations, as studies have shown that mutations can either stabilize or destabilize proteins, depending on the specific system under investigation and the methods employed for energetic analysis [4, 5].

Meanwhile, as experimental and computational techniques that can reveal conformational transitions at timescales relevant to protein function become more routine, the dynamics of proteins are increasingly recognized as an important regulator of protein function [6–8]. Reports of proteins with identical folds but different functions are not uncommon and are often attributed to differences in dynamics [9, 10]. This understanding has paved the way for enzyme engineering aimed at altering the dynamics of candidate proteins to improve catalytic efficacy [10]. In addition to highlighting the importance of dynamics in maintaining normal protein function, large-scale bioinformatics studies have suggested that DCMM sites, compared to normal residue sites, generally have tighter allosteric coupling to the functional region of the protein [11, 12], implying that DCMMs tend to alter protein function through dynamics. Although the importance of dynamics in mediating the effects of DCMMs on protein function is increasingly recognized [13, 14], how these dynamic changes affect protein-protein interactions (PPIs) remains underexplored, given that many proteins exert their functions via PPIs. A clear understanding of how disease-causing missense mutations alter protein-protein interactions, along with an in-depth exploration of the underlying mechanisms, is invaluable.

Menin is a 610-residue scaffold protein that regulates cell homeostasis in various endocrine organs by interacting with multiple proteins [15]. It is also a validated leukemia therapeutic target [16–18], against which small-molecule inhibitors are actively being developed [19]. Menin is mutation-prone, and such mutations are implicated in disease progression. It is mutated in patients with MEN1 syndrome [20, 21], as well as in sporadic parathyroid [22] and pancreatic endocrine tumors [23, 24]. More than 1,200 germline mutations in the MEN1 gene have been identified, scattered across the entire coding region without any significant hotspots or genotype-phenotype correlations [25, 26]. These mutations occur both on the surface and within the core, including at binding pockets that affect drug-Menin binding [27]. This mutation-prone nature of Menin is also evidenced by the pathogenicity score predicted by AlphaMissense [28], which shows a broad distribution of potential disease-causing mutations across the Menin protein (Fig. 1). Given its important physiological role, in silico studies exploring the effects of mutations on Menin have been reported [29, 30], but they are generally limited to studying the effects on protein stability based on crystal structures of Menin. These studies rely on Menin’s crystal structures, which may mask certain effects due to the differences between crystal and solution structures, caused by crystal packing effects. Moreover, the effects of mutations on Menin’s dynamics, and thereby on its binding with target proteins, have not been explored

**Figure 1:**
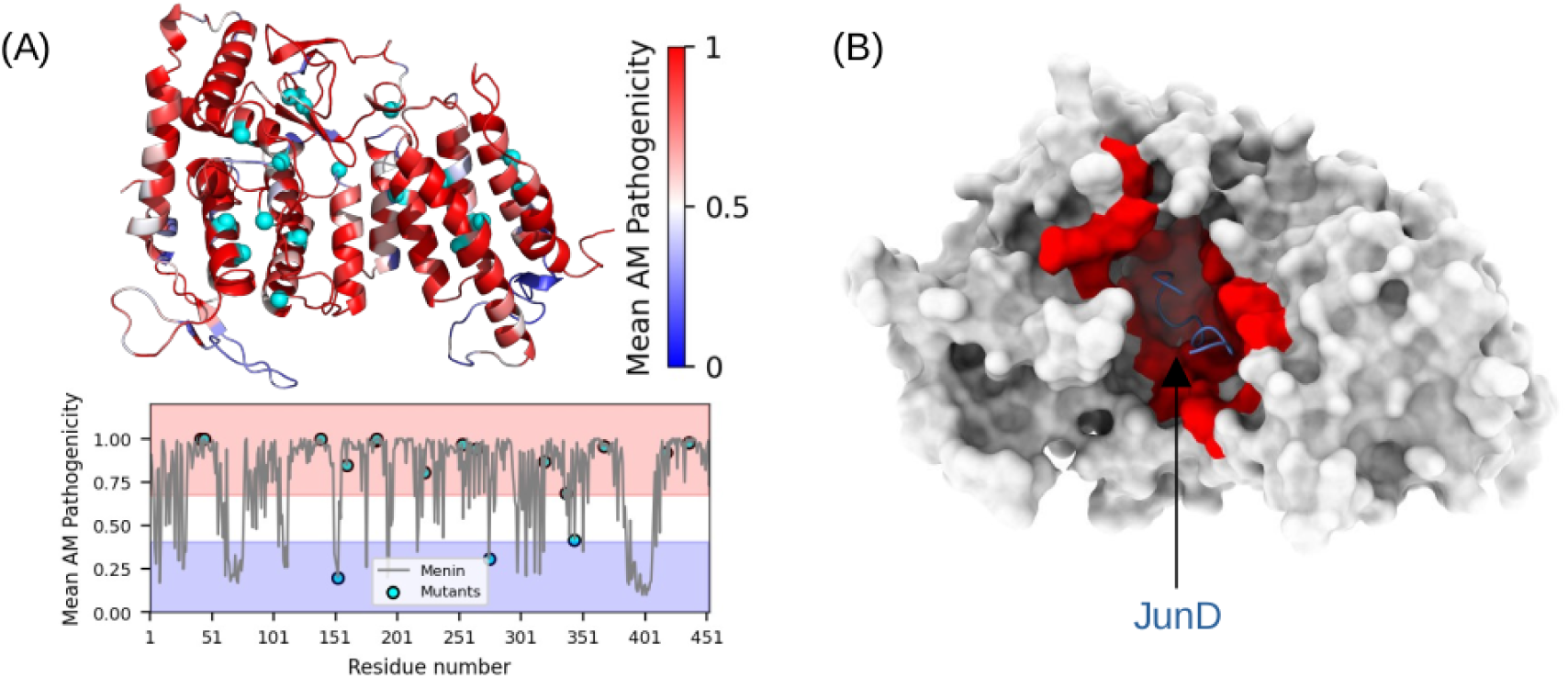
Menin’s mutation-prone nature and structural background. (A) The AlphaMissense [28] predicted pathogenicity score mapped onto the Menin protein crystal structure (PDB 3U86) [31]. A score range of 0-0.34 indicates a benign mutation, 0.34-0.56 indicates an uncertain mutation, and >0.56 indicates a pathogenic mutation. The 24 confirmed pathogenic mutations collected from the ClinVar database are shown as cyan spheres. (B) Menin is a scaffold protein with a well-organized pocket (colored red) that binds to various targets. In this study, we explore how the 24 DCMMs affect Menin’s binding with its target, JunD.

In this study, we collected 24 clinically verified pathogenic missense mutations of Menin from the ClinVar database (https://www.ncbi.nlm.nih.gov/clinvar/) (Fig. 1) and performed extensive atomistic molecular dynamics (MD) simulations on these mutants to gain insights into how they affect Menin’s protein-protein interaction (PPI) with a classic target, JunD (Fig. 1B). While we did not observe appreciable changes in protein stability induced by DCMMs, we demonstrated that they induced similar changes in protein dynamics and uniformly attenuated the Menin-JunD PPI. We further investigated the allosteric regulation of DCMMs on Menin and unraveled the mechanisms by which DCMMs attenuate the PPI. We provided a molecular picture of how DCMMs alter the Menin-JunD PPI by changing the protein’s dynamics, highlighting the necessity of explaining the effects of DCMMs on proteins through the dynamics perspective.

## 2 Methods

### 2.1 Collection of Menin mutants

We collected 24 clinically validated pathogenic missense variants of Menin from the ClinVar database (https://www.ncbi.nlm.nih.gov/clinvar/, accessed on 1st December 2022). Searching was conducted on the MEN1 gene and a total of 2,599 MEN1 mutations were recorded in the database. We included only the pathogenic mutations, excluding condition of benign, likely benign, uncertain significance, and conflicting classifications of pathogenicity, resulting in the following 24 mutants: G42A, E45G, E45K, E45Q, H139D, H139R, H139Y, D153V, A160P, W183C, V184E, L223P, S253W, L264P, R275K, P320R, A377D, T344R, A368D, D418H, D418N, D418Y, W436C and W436R. The locations of these mutations in Menin protein is shown in Fig. 1A.

### 2.2 Molecular dynamics simulations

We firstly simulated the WT and mutant Menin in the isolated state (without target). The starting structure for MD simulations was based on PDB 3U86 [31]. In this structure, residues 386-401 are missing, and therefore, we added the missing residues using a two-step strategy: 1) we simulated the fragment corresponding to the missing part in isolation, and 2) we fused the modeled fragment to the PDB 3U86 in Pymol. The Mutagenesis plugin in Pymol was used to introduce pathogenic missence mutations. The structures were first solvated into a OPC water box filled with 0.15 M KCl salts. The distance from protein to the water box wall is 12 Å, and The protein is described using the ff19SB [32] force field. The TLEAP program from Amber20 package [33] was used to generate the topology and coordinate files for MD simulations. Using the PMEMD module of AMBER20 [33], The system was first subject to 200 steps of steepest descent algorithm followed by 19800 steps of conjugate gradients algorithm for energy minimization. After minimization, the system was heated from 0 to 300 K using a two-step protocol: from 0 to 100 K over 0.04 ns in the NVT ensemble, and then from 100 to 300 K over 0.2 ns in the NPT ensemble.During the heating process, a harmonic constraint with a force constant of 5 kcal/mol/Å^2^ was applied to the protein backbone atoms. Next, a 0.4 ns equilibrium simulation was performed in the NPT ensemble with reduced force constant of 1 kcal/mol/Å^2^ at 300 K. After equilibration, each system was subjected to an independent 500 ns long production run at 300 K in the NPT ensemble. During simulations a time step of 2 fs was used, and bond lengths of all covalent hydrogen bonds are constrained using the SHAKE algorithm[34]. The non-bonded interaction cutoff was set as 10.0 Å, and longrange electrostatics were calculated using the particle mesh Particle-Mesh Ewald (PME) method.

During the MD simulations, the Langevin thermostat [35] and the Berendsen barostat were used for temperature and pressure control, respectively. The MD simulation was conducted via the AMBER20 [33] package.

We then simulated the Menin-JunD complex. To build the complex structures of mutant Menin with JunD, the last frame of the 500 ns MD of isolated Menin was extracted as starting structure. This structure was aligned with PDB 3U86 [31] on Menin via Pymol, and JunD structure from PDB 3U86 was copied and pasted into the MD-refined structure of Menin. Follow the above described MD protocol, the Menin-JunD complex was simulated for 100 ns production run.

### 2.3 Analysis of conventional MD simulations

The Rosetta-based conformational energy calculations of Menin were performed using both the crystal structure and the MD-generated structures. The crystal structures refer to Menin structure derived from PDB 3U86 (WT) and mutant structures generated via the mutagenesis plugin of Pymol based on PDB 3U86. The MD-generated structures are the last frames of 500 ns MD trajectories with waters and ions removed. These structures were first subject to refinement using the relax protocol of Rosetta prior to scoring. During the relaxation, constraints were introduced on protein backbone, leaving the sides chains to freely move. Scoring was done on the refined structures using the default energy function of Rosetta which is the REF15 [36] since July 2017. The detailed command for Rosetta scoring is listed in the supplemental file.

Analyses such as RMSD, RMSF, RoG calculations were performed using the CPPTRAJ [37] program of Amber20 package. For RMSF calculations, the MD trajectory was first aligned to the initial structure on backbone atoms. Molecular Mechanics/Generalized Born Surface Area (MM/GBSA) method was used to calculate the binding free energy (ΔG) between of Menin and JunD. Energies of the Menin-JunD complex (G*_complex_*), the isolated Menin (G*_Menin_*) and isolated JunD (G*_JunD_*) were calculated using the MMGBSA.py script from AMBER20 [33] based on the trajectory of 100 ns MD of the complex, using the modified generalized born model from Onufriev *et al* [38] with a salt concentration of 0.15 M:

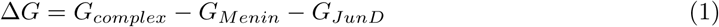

The reported ΔG is the average of 500 frames from the 100 ns trajectory, and the standard error of mean values of ΔG was reported as error bar.

### 2.4 Umbrella samplings of the Menin-JunD complex

Umbrella samplings were used to estimate the potential of mean force (PMF) along dissociation process of JunD from Menin, starting from the restart file of the 100 ns production MD run of the Menin-JunD complex. The reaction coordinate (RC) was defined as the distance between the center of mass (COM) of JunD with the COM of key Menin residues that interact with JunD in PDB 3U86. These Menin residues are L177, A182, F238, A242, C241, Y276, M278, Y319, Y323, E366 and D370. We simulated the RC range from 3 to ∼53 Å that covers the bound state to completely unbound state. The RC range was binned into 0.5 Å-wide windows, resulting in about 100 windows for each case. Harmonic restraint of 10 kcal/mol/Å^2^ force constant was introduced on the RC to force samplings at the window center. The starting structure used in each window is the last frame of the previous window. For each window, a 5 ns long sampling was conducted in the NPT ensemble at 300 K, resulting in total 500 ns sampling for each case. The PMF was estimated by using the WHAM program [39]. To get better statistics, for each case, five independent replica of umbrella samplings were conducted, and the average PMF curve was reported with error bars estimated as the standard deviation of the five PMF values at each RC point. Umbrella samplings were also performed on WT and mutant (H139D, H139R, A337D and D418N) Menin-JunD complex in which harmonic restraints of 20 kcal/mol/Å^2^ were introduced between E179 and Menin’s binding pocket. Five independent replicas of umbrella sampling runs were conducted in the presence of E179 restraints. All MD simulations performed in this study were summarized in Table S1.

### 2.5 Dynamic coupling index (DCI) analysis

The DCI analysis is a method that quantifies the strength of allostery coupling between residue pairs, usually between mutation sites and function important regions of the protein [11, 14]. Briefly, this method firstly records the conformational displacement of a protein in response to random perturbation on residues. The displacement is calculated using the linear response theory (LRT) theory:

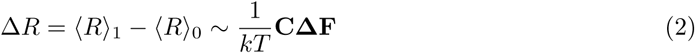

where ΔR is the conformational displacement, ΔF is the random perturbation force acting on residue, and **C** is the variance-covariance matrix. For a protein that consists of N residues, the conformational displacement can be written as the following matrix [14, 40]:

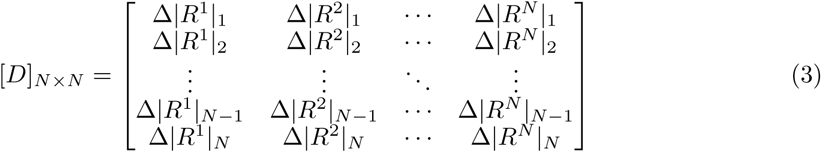

where |ΔR*^j^*|*_i_* = ⟨(ΔR)^2^⟩ is the displacement of residue i in response to the perturbation at residue j. To shed insights into how mutation site could impact the protein function through allostery regulation, one can first designate an functional import regions in a protein (usually the catalytic site or binding pocket) and record the displacement of mutation site in response to perturbation of the functional region. Following this idea, the DCI score of a residue i is defined as follow [11]:

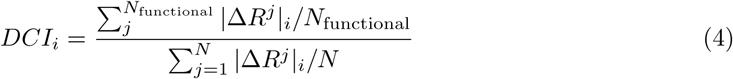

where N*_functional_* is the total number of residues that constitute the functional region of the protein. The denominator is the total displacement of residue i in response to perturbation at all residues in the protein. In our case, these functional residues are those form the binding pocket of Menin, namely, the residues that are within 3.5 Å of JunD in the PDB 3U86. The covariance matrix (inverse Hessian) used in LRT for conformational displacement estimation was calculated via an elastic network model (ENM) on the last frame of 500 ns MD trajectory. The python scripts used for DCI analysis were adapted from Modi *et al* ’s work [10] (https://github.com/SBOZKAN/DFI-DCI).

## 3 Results

### 3.1 Pathogenic mutants do not necessarily destabilize protein stability

We first evaluated the effects of DCMMs on Menin’s structural stability by calculating conformational energy. A previous study used FoldX [41] to calculate the folding energy difference between mutants and WT Menin (ΔΔG*_conf_*) based on crystal structures and reported that DCMMs tend to exhibit larger positive ΔΔG*_conf_* values compared to benign mutations [29]. This finding suggests that disease-causing mutations are more likely to destabilize Menin’s structural stability. Here, we re-examine the impact of pathogenic mutations on Menin’s conformational stability by scoring the protein structure using the REF15 Rosetta energy function [36]. The REF15 energy function is highly accurate in ranking native protein conformations as the lowest-energy states [42, 43]. Our approach is further motivated by studies showing that different computational methods for calculating conformational energy can yield varying results, and combining scores from different methods generally improves accuracy [44].

We present the Rosetta-based ΔΔG*_conf_* values for mutants relative to WT Menin in Fig. 2A. Compared to WT Menin, which has a conformational energy of −928.877 kcal/mol, the 24 mutants studied exhibit either positive or negative ΔΔG*_conf_* values. Using |ΔΔG*_conf_* | > 10 kcal/mol as the threshold for significant structural impact, we found that 54.7% of the mutants show negligible effects on Menin’s stability. Among the remaining 45.3%, six mutants (E45G, D153V, A160P, W183C, L223P, and L264P) exhibit positive ΔΔG*_conf_* values, indicating destabilization. Notably, A160P, L223P, and L264P are particularly destabilizing, with |ΔΔG*_conf_* | > 10 kcal/mol. Conversely, five mutants (E45Q, H139Y, P320R, T344R, and D418) exhibit negative ΔΔG*_conf_* values below −10 kcal/mol, suggesting stabilization of Menin’s structure. Overall, the Rosetta energy scores based on crystal structures suggest that disease-causing mutations can either stabilize or destabilize Menin’s structure, with destabilizing effects slightly more prevalent but within the margin of error. To further validate these findings, we repeated the conformational energy analysis using Menin structures obtained from the final frames of 500 ns MD simulations (Fig. 2B). While individual ΔΔG*_conf_* values varied significantly after MD refinement, the overall trend remained consistent: the 24 mutants either increased or decreased Menin’s conformational energy, exhibiting a non-uniform impact on structural stability. Hydrogen bonding analysis showed that mutants that increased or decreased ΔΔG*_conf_* values relative to WT Menin tended to lose or gain hydrogen bonds (Fig. S1), respectively, further corroborating their stabilizing or destabilizing effects.

**Figure 2:**
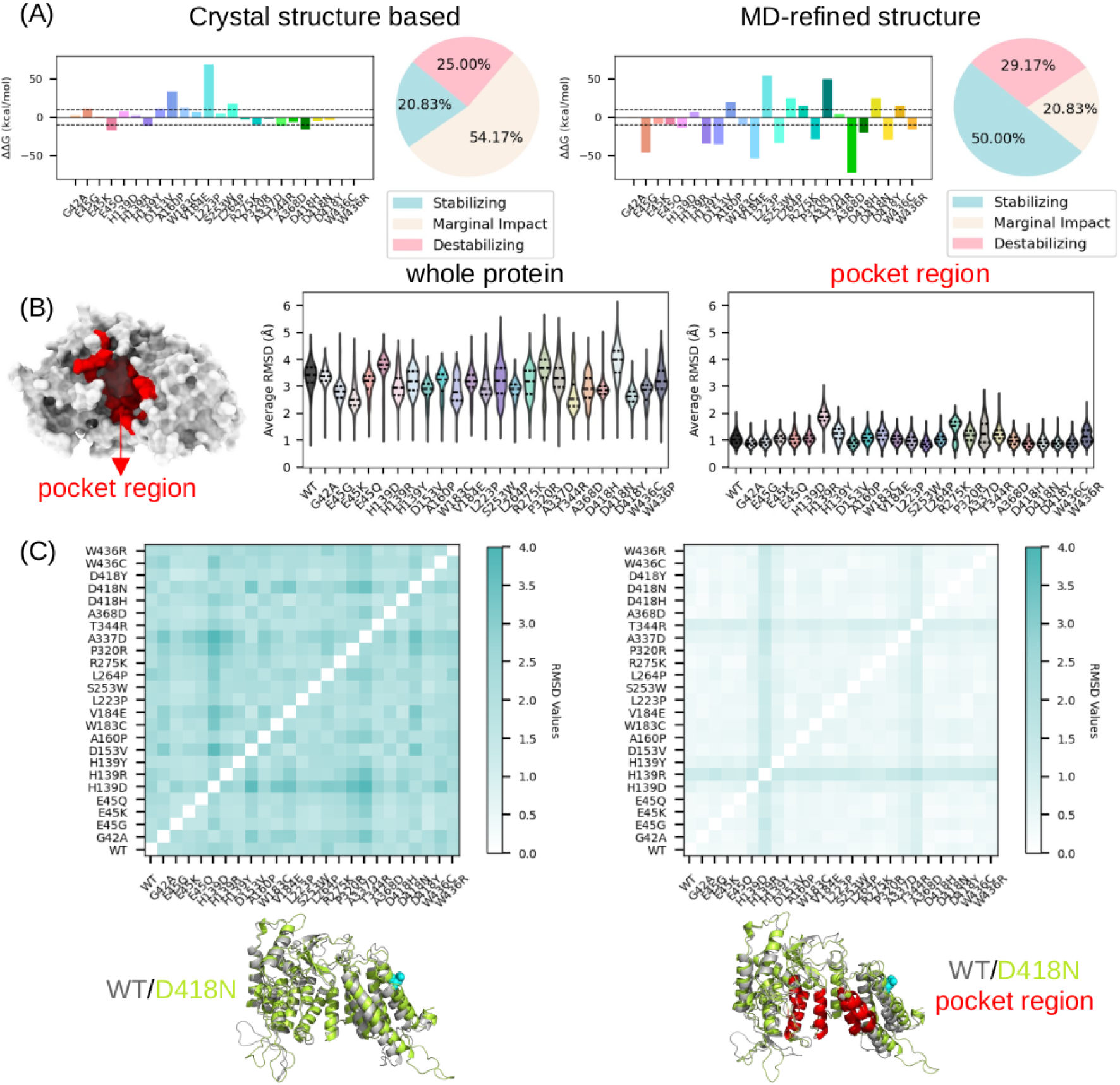
Effects of mutations on Menin protein stability. (A) Conformational energy of the Menin protein calculated using the REF15 Rosetta function [36] based on crystal structure and 500 ns MD-refined structures. The conformational energy difference of the mutant relative to WT Menin (ΔΔG*_conf_*) is reported. The two dashed lines represent ΔΔG*_conf_* = 10 and −10 kcal/mol, respectively. Mutants with |ΔΔG*_conf_* | ≤ 10 are considered to have a marginal impact on protein stability, while those with ΔΔG*_conf_* > 10 and < −10 kcal/mol are considered destabilizing and stabilizing mutations, respectively. (B) Average RMSD of backbone atoms of the whole Menin protein and of the Menin pocket (defined as residues within 3.5 Å of JunD in PDB 3U86). RMSDs were calculated relative to the starting structures of the MD simulations. (C) Pairwise RMSD between Menin mutants calculated using the average structure from the MD simulation. Overlay of MD-refined WT Menin and the mutant D418N.

We further analyzed the effects of pathogenic mutants on the structural properties of Menin by comparing the root mean squared deviations (RMSD) between WT Menin and mutants based on the 500 ns MD trajectory. Consistent with the Rosetta energy scores, we found that during the MD simulation, both WT and mutants underwent comparable conformational changes relative to the crystal structure 3U86 (Fig. 2C). The pairwise RMSD matrix, calculated using the average structure from the 500 ns MD simulations, clearly shows that WT and mutants exhibit very similar conformations, with most RMSD values around 2 Å. We also show the MD-refined structures of WT Menin and the mutant D418N, which exhibits the largest RMSD of 3.91 Årelative to PDB 3U86. Despite this, we observed no significant conformational changes in the overall fold of Menin. A zoomed-in view of the functional pocket reveals that the D418N mutant maintains a nearly identical pocket structure to WT Menin, with an RMSD of around 1 Å. Measurements based on the radius of gyration and surface area (Fig. S1) are consistent with the RMSD analysis, further demonstrating that proteins bearing disease-causing mutations undergo conformational changes comparable to those of WT Menin. **Based on extensive all-atom MD simulations, we show that Menin mutations do not necessarily destabilize the conformational stability of Menin.**

### 3.2 Pathogenic mutants cause similar changes in protein dynamics

To investigate the effects of missense mutations on the dynamics of the Menin protein, we calculated the root mean squared fluctuations (RMSF) of the protein using the 500 ns MD trajectory and compared the per-residue RMSF differences (ΔRMSF, mutant minus WT). As shown in Fig. 3A, plotting the ΔRMSF curves revealed that mutants caused similar changes in the dynamics of the Menin protein, as evidenced by uniformly increased or decreased RMSF values in several regions. To assess the similarity of dynamic changes across mutants, we analyzed whether these mutants, when represented as points in a higher-dimensional space using their per-residue ΔRMSF values as coordinates, could be grouped into separate clusters. Our rationale is that if the mutants can be well grouped, the dynamic changes are heterogeneous; otherwise, the dynamic changes are similar across the mutants. To this end, each Menin mutant was encoded as an N-dimensional vector, ΔRM⃗SF*_N_* = (ΔRMSF_1_, ΔRMSF_2_, .., ΔRMSF*_N_*), where N is the number of residues in Menin. These high-dimensional points were then projected onto a 2D plane after principal component analysis (PCA), and K-means clustering was performed on the projected data. As shown in Fig. 3B, the Silhouette scores, which quantify the quality of data separation in clustering, were all ≤ 0.45 across different cluster numbers, indicating that the mutants cannot be well grouped into distinct clusters. Even for a cluster number of four, which gave the highest Silhouette score (0.45), the resultant clusters were still relatively mixed, with only a few outliers. Therefore, the clustering analysis supports the conclusion that these Menin mutants induce similar RMSF changes in Menin (Fig. 3B).

**Figure 3:**
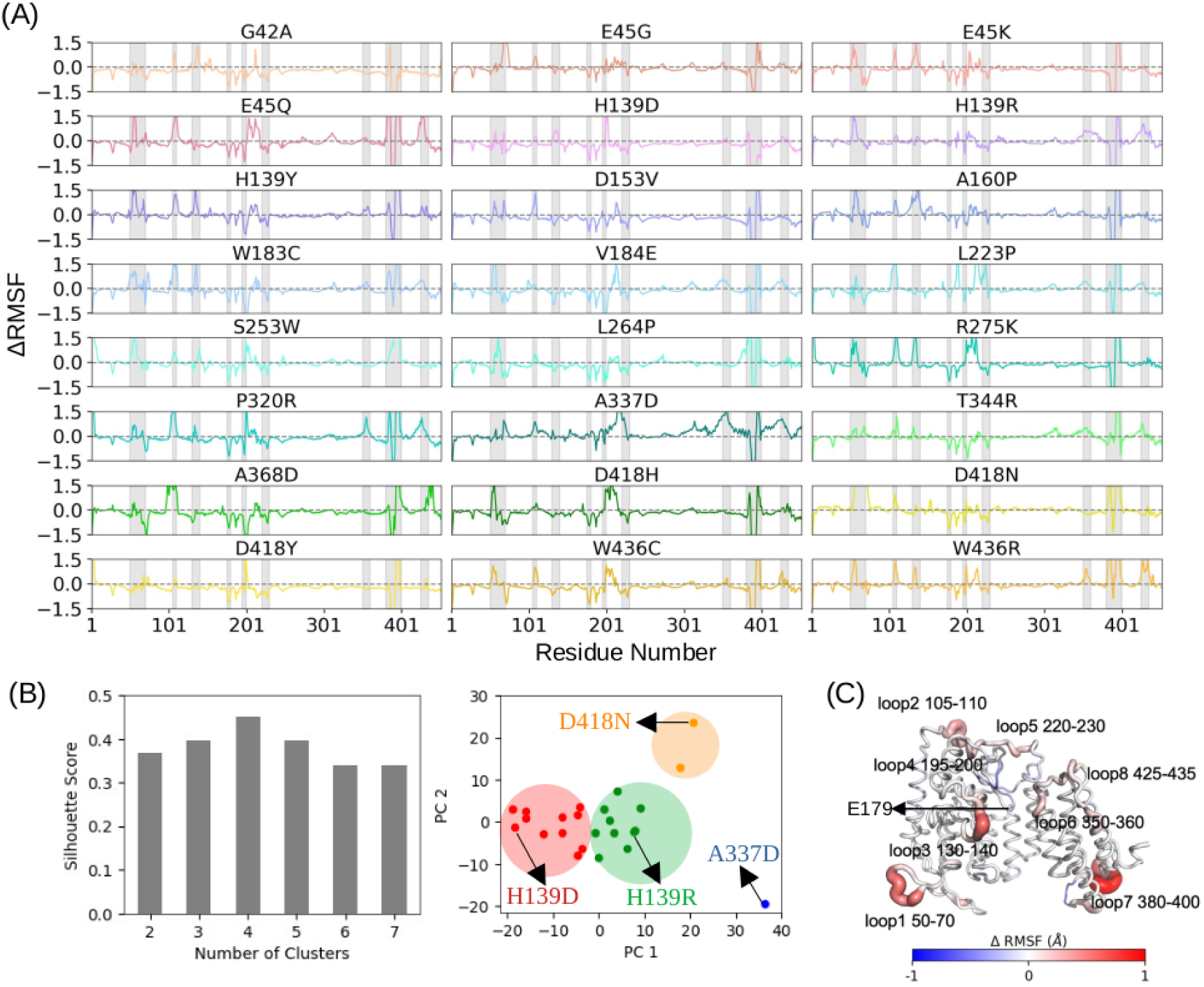
**Effects of mutations on Menin protein dynamics**. A) Per residue ΔRMSF (mutant minus WT) curves. (B) Silhouette scores of K-means clustering on mutants. Mutants were encoded as high dimensional vectors using their per-residue ΔRMSF values. The clustering results at four cluster number were shown. (C) Mapping of the average ΔRMSF over all mutants on Menin structure (PDB 3U86), with eight regions that have large RMSF difference labeled. These regions are also highlighted in panel A.

To identify the regions where dynamics are significantly altered by mutations, we calculated the average per-residue ΔRMSF across all mutants and mapped the values onto the Menin crystal structure (PDB 3U86) (Fig. 3C). We highlighted eight regions: loop 1 (residues 50-70), loop 2 (residues 105-110), loop 3 (residues 130-140), loop 4 (residues 195-200), loop 5 (residues 220-230), loop 6 (residues 350-360), loop 7 (residues 380-400), and loop 8 (residues 425-435), where the dynamic changes are most pronounced. Interestingly, although the dominant effect of the mutants is to increase the flexibility of peripheral loops, motifs adjacent to the binding site are also affected. These motifs include loop 4 and loop 8, whose dynamics are respectively attenuated and accentuated, suggesting that these dynamic changes could have consequences for Menin’s interactions with its targets. We showed that all Menin pathogenic mutants alter the dynamics of the Menin protein compared to the WT protein, and these dynamic changes are similar across the mutants.

### 3.3 Pathogenic mutations attenuate the Menin-JunD PPI

To explore the effects of pathogenic mutants on the protein-protein interaction (PPI) of Menin with its targets, we selected JunD, a well-characterized Menin target that regulates cell proliferation [45, 46] after binding to Menin. Specifically, we performed umbrella samplings to simulate the dissociation of JunD from Menin, estimating the potential of mean force (PMF) along the process and uncovering the dissociation pathways. We selected WT Menin and four representative Menin mutants (each from a different group clustered in Fig. 3B) for constructing the Menin-JunD complex, which was subsequently subjected to umbrella samplings for dissociation studies. To ensure sufficient sampling for better PMF statistics, we performed five independent replicas of umbrella samplings for each case.

In Fig. 4A, we show the average PMF curves along the dissociation process. By defining the bound state and unbound states at RC = 7.6 and 50 Å, respectively, the Menin-JunD binding free energy (ΔG) for WT is −53.18 ± 8.75 kcal/mol. More importantly, we found that the Menin-JunD ΔG in the mutants is significantly larger than that of WT Menin, with an average ΔΔG ∼27 kcal/mol in favor of the WT-JunD binding. This suggests that, compared to WT Menin, the mutants have reduced binding affinities for JunD. We also observed that the reduced binding affinity in mutants is mainly attributed to increased unbinding rates. Based on the PMF curves, the energy barrier JunD needs to overcome before completely leaving the Menin pocket is higher in WT than in the mutants (Fig. 4A). The height of the barrier, and thus the slope of the free energy curve from bound state to transition state, is closely related to the unbinding rate (k*_off_*) [47]. Therefore, JunD in the mutants exhibits faster unbinding rates compared to WT.

**Figure 4:**
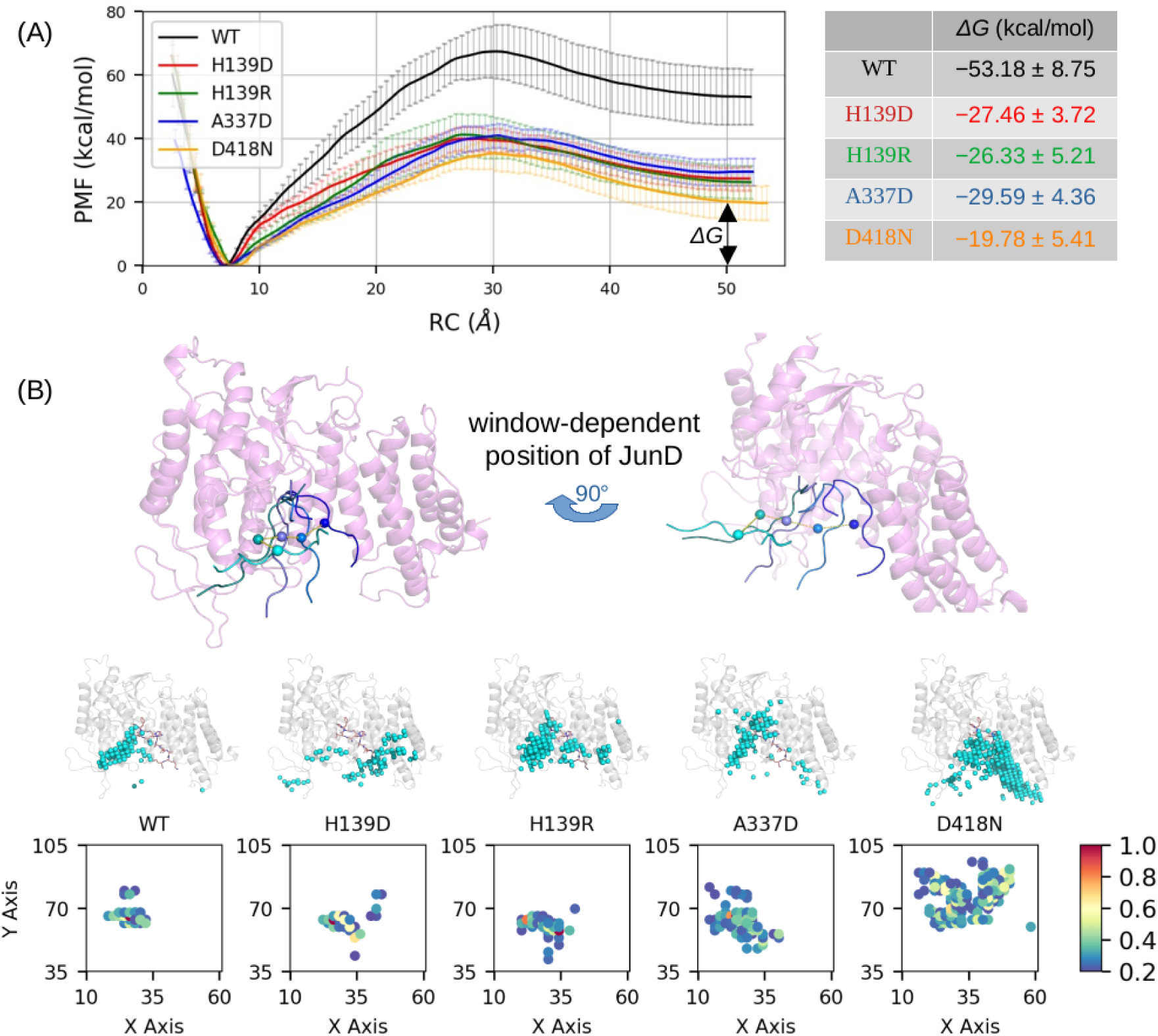
Effects of mutations on Menin-JunD PPI. (A) PMF curve along the dissociation process of JunD from Menin. The average PMF curve from five replicas of umbrella sampling is shown, with error bars representing the standard deviation of PMF values at each RC point. The free energy difference (ΔG) was calculated as ΔG = PMF*_bound_* − PMF*_unbound_*, where the bound and unbound states correspond to RC values of 7.6 and 50 Å, respectively. (B) The window-dependent positions of JunD were extracted from umbrella sampling to outline the dissociation pathway. The distribution of the COM of JunD is shown below, with each dot representing a grid point where the COM density is >0.2. The projection of these grid points onto the X-Y plane is visualized, with density values indicated by a color bar.

The altered unbinding kinetics of JunD are also reflected in the dissociation pathways of the mutants relative to WT Menin. We extracted the window-dependent positions of JunD from the umbrella samplings and plotted the center of mass (COM) of JunD to delineate the dissociation pathways (Fig. 4B). We found that in WT Menin, JunD follows a highly conserved dissociation pathway across the five umbrella samplings (Fig. S4). In WT Menin, JunD follows a path mainly formed by Menin residues 130-140 and 240-255 during unbinding, while such conserved pathways are disrupted in the mutants. Specifically, JunD has diminished contacts with Menin residues 130-140 and 240-255 in the mutants but gains contacts with other regions, such as residues 350-400 and 430-440, during unbinding (Fig. S2). To better illustrate the dissociation pathways, we plotted the COMs of JunD sampled during the accumulated umbrella samplings and projected them onto a 2D plane (Fig. 4B). We demonstrated that JunD in Menin mutants follows diverse dissociation pathways, as evidenced by the spreading of JunD COMs over Menin’s pocket, in contrast to the concentrated, strip-like COM distributions observed in WT Menin (Fig. 4B). **Overall, we demonstrated that disease-causing mutations attenuate the Menin-JunD PPI and disrupt the highly conserved dissociation pathway observed in WT Menin.**

### 3.4 Pathogenic mutants disrupt the coupling between E179 and JunD binding site through allostery

In the PMF calculations, the mutations that attenuate the Menin-JunD PPI are located far from Menin’s binding pocket, with an average distance of 21.8 Å. To investigate the mechanism by which these mutations weaken the Menin-JunD interaction, we examined their allosteric regulation of Menin’s function. The dynamic coupling index (DCI) analysis, introduced by Özkan et al. [11], quantifies the allosteric coupling strength between arbitrary residues and designated functional regions (typically functionally important residues) in a protein. Higher DCI values indicate stronger couplings. This method has proven effective in elucidating the allosteric effects of mutations on protein function [11] and has guided enzyme engineering by modifying protein dynamics to enhance catalytic efficiency [10]. Here, we employed DCI analysis to calculate the allosteric coupling between Menin residues and the JunD binding pocket (functional site). As shown in Fig. 5A, compared to normal residue sites, mutations are generally located farther from the functional site yet exhibit stronger allosteric couplings with it. This observation suggests that Menin mutations predominantly affect protein function through allostery. Furthermore, it aligns with previous DCI-based studies demonstrating that allostery is a primary mechanism by which disease-causing missense mutations alter protein function [11, 12].

**Figure 5:**
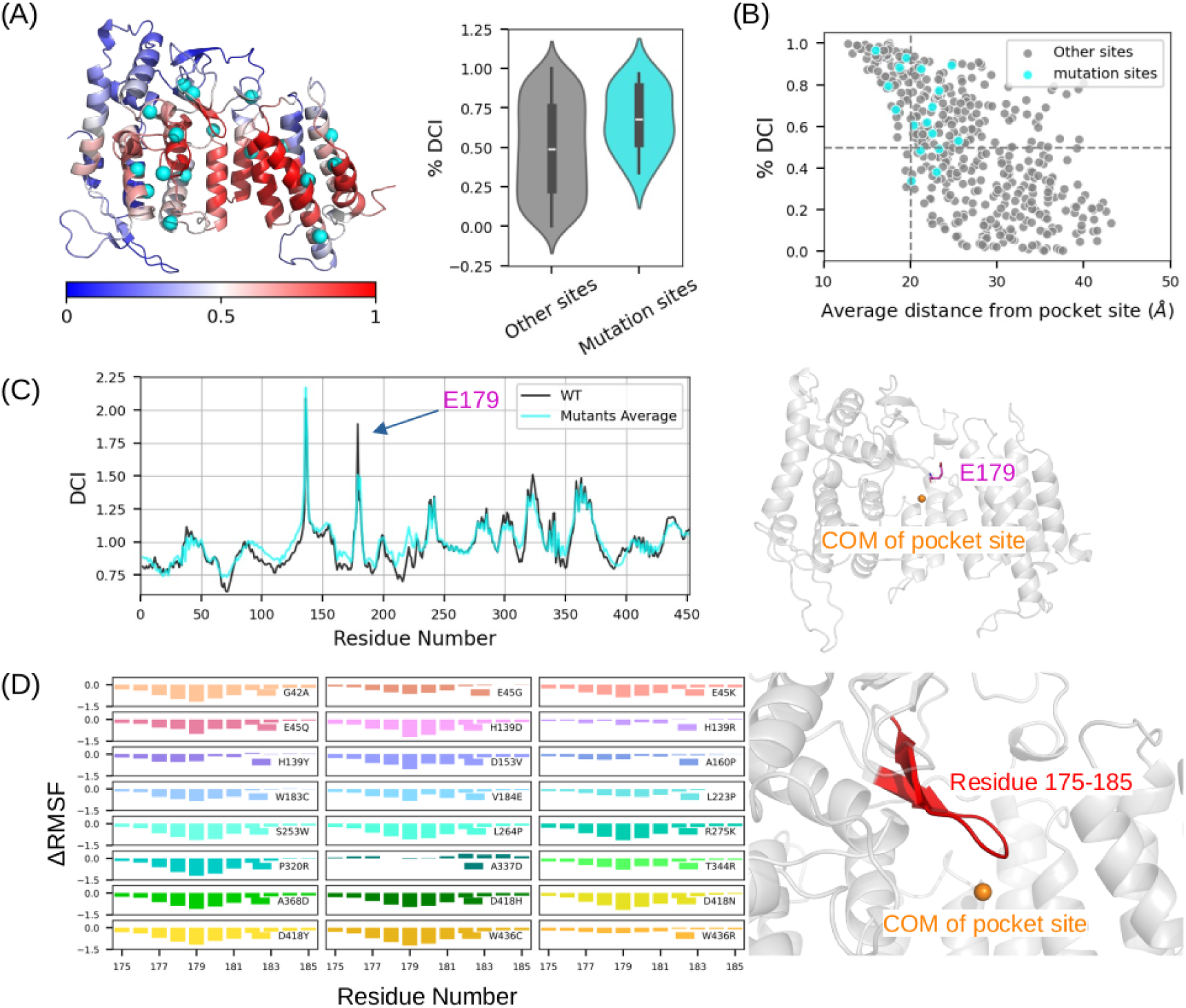
Allosteric Regulation Effects of Mutations. (A) Ribbon diagram showing the %DCI as a color-coded spectrum from blue to white to red, representing weak to strong allosteric coupling to the functional site. Mutant residues are displayed as cyan spheres. The distribution of DCI values for normal and mutant sites is shown alongside. (B) Scatter plot of all Menin residues based on their %DCI values and distances from the functional site. Dashed lines mark the thresholds at %DCI = 0.5 and distance = 20 Å. (C) Average per-residue DCI values for mutants compared to WT Menin. Residue E179 shows a reduced DCI value in mutants relative to WT. The position of E179 relative to the COM of the functional pocket is shown on the right. (E) ΔRMSF values for residues around E179 (mutants minus WT). The corresponding fragment’s location in the Menin structure is shown on the right.

To link the tighter allosteric coupling in Menin mutants to the impaired JunD binding, we compared the per-residue DCI value differences between mutant and WT Menin, aiming to identify specific motif(s) responsible for the PPI alterations. As shown in Fig. 5C, we plotted the DCI curve of WT Menin alongside the average DCI curve over all 24 mutants. Interestingly, residue E179 exhibited a notable reduction in its DCI value, dropping from 1.90 in WT Menin to 1.51 in the mutants. This indicates a significant weakening of the coupling between E179 and the binding pocket in the mutants. Further analysis of the individual DCI curves for each mutant confirmed this trend: 22 out of the 24 mutants (with the exceptions of L223P and R275K) showed a reduced DCI value for E179 (Fig. S5). Moreover, we observed that the mutants uniformly rigidified the region around E179, as evidenced by the reduced RMSF values for residues 175–185 compared to WT Menin (Fig. 5D). The diminished dynamics of this region likely contribute to the decoupling of E179 from the functional pocket. In Fig. 4, we identified a conserved JunD dissociation pathway in WT Menin, where residue E179 maintains contacts with JunD during the early stages of unbinding. We speculate that, in mutants, the reduced coupling between E179 and Menin’s pocket reshapes the interaction energy landscape near the bound state, thereby weakening the Menin-JunD interaction. **Based on DCI analysis, we demonstrated that DCMMs, despite being located far from the binding pocket, are tightly coupled to Menin’s functional region. Furthermore, a consistent reduction in the coupling between E179 and the functional site was observed across nearly all mutants, likely contributing to the attenuated Menin-JunD PPI.**

### 3.5 Attenuated PPIs in mutants can be remedied by forcibly coupling E179 with the JunD binding site

We identified that the coupling between E179 and Menin’s functional pocket was disrupted in the mutants, likely contributing to the attenuated Menin-JunD PPI observed in these systems. To test whether restoring this coupling could strengthen the Menin-JunD interaction, we introduced harmonic restraints between E179 and the center of mass (COM) of the functional pocket as a means of forcibly maintaining the coupling (Fig. 6A). Umbrella sampling simulations were then performed to study the effects on the PPI. As shown in Fig. 6A, a comparison of the average PMF curves in the presence and absence of E179 restraints reveals that, for all mutants, introducing the E179 restraint significantly lowered the Menin-JunD binding free energy (ΔG) by approximately 10–20 kcal/mol relative to the unrestrained condition, demonstrating enhanced Menin-JunD binding affinity. Interestingly, in the WT system—which already exhibits optimal E179–functional site coupling—the ΔG was only slightly affected by the E179 restraint, changing from −53.18 kcal/mol to −49.81 kcal/mol. This result suggests that the E179 restraint has a negligible impact on the Menin-JunD PPI in WT systems. Beyond binding free energy, we also examined the dissociation pathways in the presence of the E179 restraint. For the mutants, forcibly coupling E179 to the functional pocket led to less diverse JunD dissociation pathways compared to those observed previously (Fig. 4B). Specifically, with E179 restraints in place, the 2D projections of JunD COMs during the umbrella sampling simulations displayed more concentrated distributions. This behavior increasingly resembled the conserved dissociation pathway observed in WT Menin, where the dissociation pathway remains largely unaffected by E179 restraint. **Thus, for all four mutants, forcibly maintaining the disrupted coupling between E179 and the functional pocket rescues the attenuated Menin-JunD interaction. These findings** confirm that the detrimental effects of the mutants on Menin-JunD PPI are exerted through allosteric disruption of the E179–functional pocket coupling.

**Figure 6:**
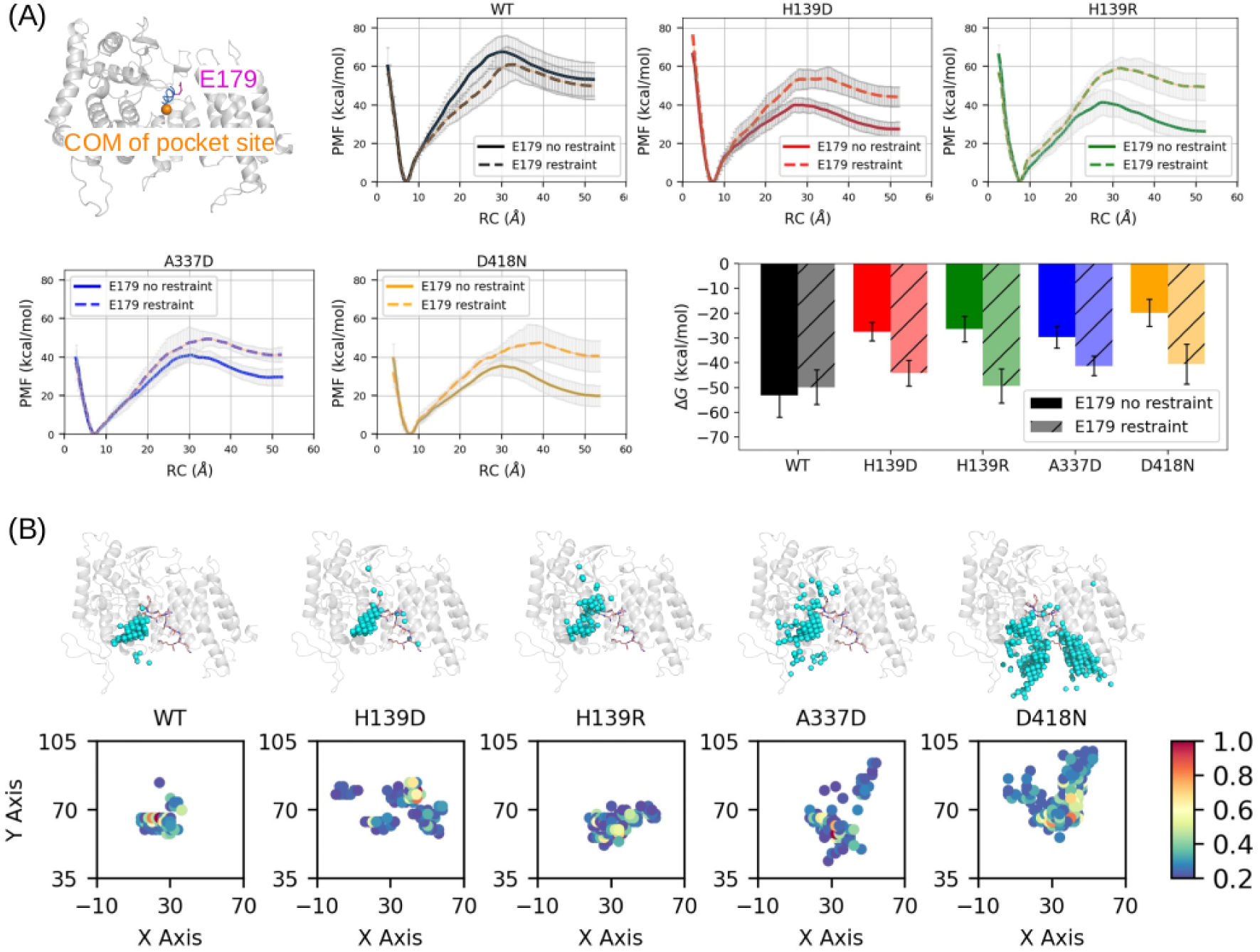
Menin-JunD PPI with forced E179-Pocket coupling. (A) Schematic illustration of the 20 kcal/mol/Å^2^ harmonic restraints applied between E179 and the center of mass (COM) of the functional pocket. The PMF curves along the Menin-JunD dissociation pathway, with and without the imposed restraint, are shown. (B) Distribution of the COM of JunD during PMF sampling. Each dot represents a grid point where the normalized density of the COM of JunD is greater than 0.2, as calculated using the ”grid” command in the CPPTRAJ program. The X and Y coordinates of these grid points are projected onto a 2D plane to visualize the COM distributions more clearly.

## 4 Discussion

Our examination of mutations’ effects on Menin protein stability is warranted. Explaining the effects of disease-causing missense mutations on protein function from a structural stability perspective is appealing, given that native proteins with evolutionarily selected amino acids are located at the bottom of the funneled conformational energy landscape [48]. It is tempting to argue that perturbation of the amino acid sequence would raise the conformational energy, thereby destabilizing the protein’s stability. However, for a protein to function, the roughness of the energy landscape surface that dictates its functional dynamics is also important [49]. Therefore, it is necessary to examine the effects of pathogenic mutations on protein dynamics, as mutations may change the roughness of the landscape while still maintaining the overall funneled free energy landscape. Specifically, for Menin, similar protein stability analyses based on conformational energy calculations on crystal structures have been reported [29, 30]. However, such analyses have limitations, such as the fact that thermodynamic calculation outcomes are methodology-dependent [5], and crystal structures may not reflect the functional state of the protein under physiological conditions. Given this, we reexamined the effects of mutations on Menin’s structural stability, emphasizing the following points in our evaluation process: 1) We used the REF15 Rosetta energy function [36], a community-recognized scoring function that is accurate in ranking the native protein structure as the lowest-energy structure [42, 43] and is also reliable in predicting mutation-induced stability changes in proteins [50]. 2) We performed 500 ns atomistic equilibrium MD simulations to relax the system and evaluate the stability using both crystal structures and MD-refined structures. 3) We selected clinically verified pathogenic mutations that are broadly distributed in Menin, including buried mutations and those near the surface of Menin. We show that, when assessed by the Rosetta energetic scores (ΔΔG*_conf_*), pathogenic missense mutations can either stabilize or destabilize the Menin protein, showing no uniform trend of destabilizing the protein. When assessed by structural metrics (e.g., RMSD), the mutants underwent comparable conformational changes to those of the WT protein during the MD simulations. These data together suggest that mutations do not necessarily destabilize Menin, and their impacts on protein function need to be scrutinized from a protein dynamics perspective.

Due to computational cost considerations, we selected the WT protein plus four representative mutants for umbrella sampling studies instead of simulating all 24 mutants. To verify the observations from umbrella sampling, which indicated that the mutants reduced the binding affinity between Menin and JunD, we performed the more efficient MMGBSA-based ΔG calculations for all mutants. Briefly, we built the Menin-JunD complex structure using the MD-refined Menin structure and the aligned JunD structure from 3U86 [31], and ran 100 ns atomistic MD simulations. We present the MMGBSA-based ΔG values for all mutants in Fig. 7A, and found that 22 out of 24 mutants exhibited reduced binding to JunD compared to WT Menin, with the most significant reductions in binding affinity (16–25 kcal/mol) observed for H139Y, D153V, A160P, W183C, and V184E. Only three Menin mutants (H139R, S253W, and R275K) maintained comparable affinities to WT Menin. These data are consistent with the umbrella sampling results, demonstrating that the mutants attenuated Menin-JunD binding affinities. Interestingly, we found that in the presence of JunD, the effects of mutations on Menin’s structural properties became more pronounced: Menin mutants now underwent larger conformational changes during the 100 ns MD simulations compared to WT Menin (see RMSD comparison in Fig. 7B and radius of gyration comparison in Fig. S3). The conformational differences between Menin mutants also became larger, as assessed by the pairwise Menin RMSD matrix calculated using the average structure from the MD simulations. Therefore, the presence of JunD sensitized Menin’s conformational stability to disease mutations, causing the mutants to undergo greater conformational changes than WT Menin While mutations do not cause severe conformational changes in the isolated Menin protein, they do lead to similar patterns of changes in protein dynamics. These dynamic changes mainly occur in the loop regions of the Menin protein, including the peripheral loops far from the binding site and loops adjacent to the site. Similar dynamic changes in loops that have functional consequences have been reported in proteins such as the green fluorescent protein family [51]. Indeed, loops are vital connector regions in proteins, and their dynamics are, to some extent, representative of the protein’s functional dynamics [52]. Therefore, the dynamic changes we observed in Menin are highly likely to be related to alterations in Menin’s function. Using extensive umbrella sampling, corroborated with large-scale MMGBSA-based ΔG binding free energy calculations, we demonstrated that mutations attenuated Menin’s binding to a classic target, JunD, revealing the consequence of these mutations on Menin’s function As a scaffold protein, Menin possesses a conserved pocket to bind various protein targets to exert its function. To understand how mutations transmit signals from a distance to the binding pocket to affect PPIs, we performed dynamic coupling index (DCI) analysis. This DCI method has been proven effective in quantifying the allosteric coupling between distant residues and the functional region of proteins [10, 53, 54]. We found that, compared to normal residue sites, the mutation sites, which are on average 21.15 Å away from Menin’s pocket, have an average DCI of 0.68%, compared to 0.49% DCI for normal sites. This finding is consistent with large-scale DCI analyses on other proteins, which show that pathogenic missense mutations, although far from the functional site, have tighter allosteric couplings with the functional site [11]. More importantly, we discovered that in all mutants, a key residue, E179, had its dynamic coupling with the pocket dramatically diminished. This finding provides a clue linking the overall tight dynamic coupling of the mutants to alterations in Menin’s function. To verify if E179 could serve as an actionable target to rescue the attenuated PPI, we performed repeated umbrella samplings in the presence of harmonic constraints that forcibly maintained the coupling between E179 and the pocket. Excitingly, we found that the Menin-JunD PPI was strengthened. Therefore, our analysis of the mutations’ allosteric effects on Menin function—ranging from whole-protein level DCI analysis to the pinpointing of a key residue responsible for the altered PPI—provides a molecular picture of how pathogenic missense mutations change Menin’s function through allostery.

**Figure 7:**
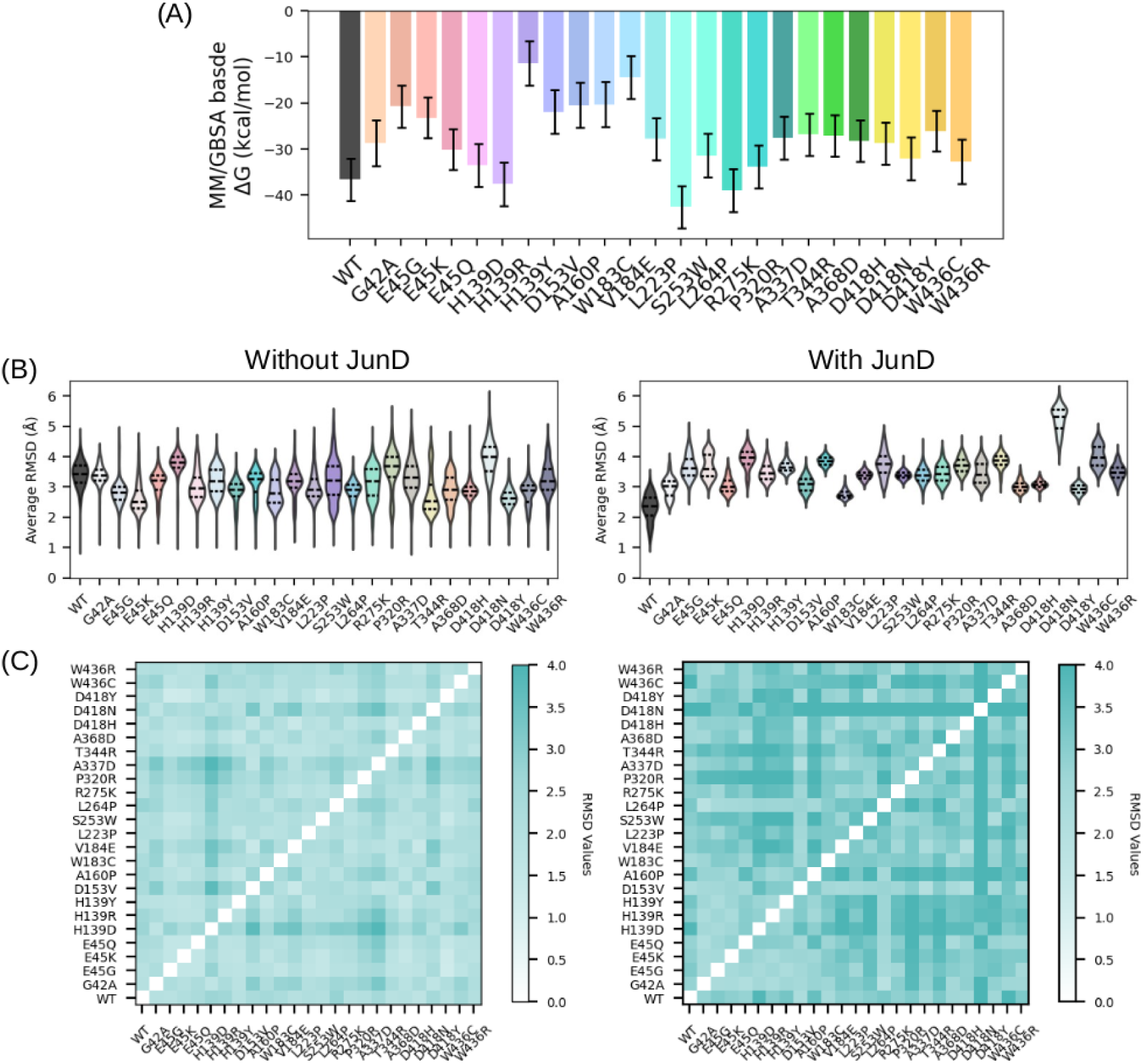
**Effects of mutations on Menin protein stability in the presence of JunD target**. (A) MMGBSA calculated binding free energy (ΔG) between JunD and Menin. (B) The average backbone RMSD of Menin relative the PDB 3U86 with and without JunD. (C) The pair-wise backbone RMSD matrix of Menin.

Our findings support the notion that dynamics is not only important for the normal function of proteins but also mediates the effects of disease-causing mutations on protein-protein interactions. Missense mutations are frequently observed in the human genome [11], and they do not necessarily cause disease because pairs of missense mutants can compensate for each other’s detrimental effects on the protein’s dynamics, keeping dynamics unchanged relative to the WT protein [12]. Our demonstration of the mutants’ allosteric effects on the protein, as well as the identification of specific motifs that can revert the detrimental effects of mutants on the PPI, exemplifies the necessity of understanding pathogenic mutations’ effects from a protein dynamics perspective rather than merely from a structural perspective. Our results support the idea that dynamics may be a more universal indicator of protein function changes and align with the ever-prevailing viewpoint that protein dynamics are indispensable in explaining the effects of pathogenic mutations on protein functions [55].

## 5 Conclusions

By using extensive conventional MD and umbrella sampling combined with allostery analysis, we show that disease-causing missense mutations alter Menin’s PPI with its target by changing the protein’s dynamics. We first demonstrated that mutations can either increase or decrease the Rosetta-scored conformational energy relative to the WT protein. This finding is consistent between conformational energy calculations using crystal structures and MD-refined structures, suggesting that mutations do not necessarily destabilize the protein. However, we found that mutations change the dynamics of the protein, and the patterns of dynamic changes are similar across the mutants. The consequence of such dynamic changes on Menin’s PPI with JunD was probed by umbrella sampling, in which we showed that mutations attenuate the PPI binding stability and alter the dissociation pathway along which JunD unbinds from Menin. The allosteric regulation mechanism of mutations on the Menin-JunD PPI was further probed by DCI analysis, where we showed that mutations are tightly coupled to Menin’s functional site, despite being far away from it. Furthermore, we identified a key residue, E179, whose coupling with Menin’s pocket is disrupted due to the allosteric effects of mutations. The vital role of E179 in the PPI was verified as we showed that forcibly maintaining E179’s coupling with Menin’s pocket rescued the impaired PPI in mutants. This suggests that E179 is an actionable target that could be used to counterbalance mutations’ effects on the Menin-JunD PPI. Overall, our data provide a molecular picture of how disease-causing missense mutations alter Menin’s function through dynamics, exemplifying the importance of dynamics in mediating the function of pathogenic protein variants.

## Acknowledgements

Research reported in this work was supported by the National Natural Science Foundation of China (22002096), China Postdoctoral Science Foundation (2023M730827), Heilongjiang Post-doctoral Science Foundation (LBH-223123), Natural Science Foundation of Heilongjiang Province (YQ2023H015), Heilongjiang Province Funding Project for Returned Overseas Scholars (21032220010) and Harbin Medical University high-level introduction of talent research start-up funds (31011210004 and 31021200109).

---

**Figure S1:**
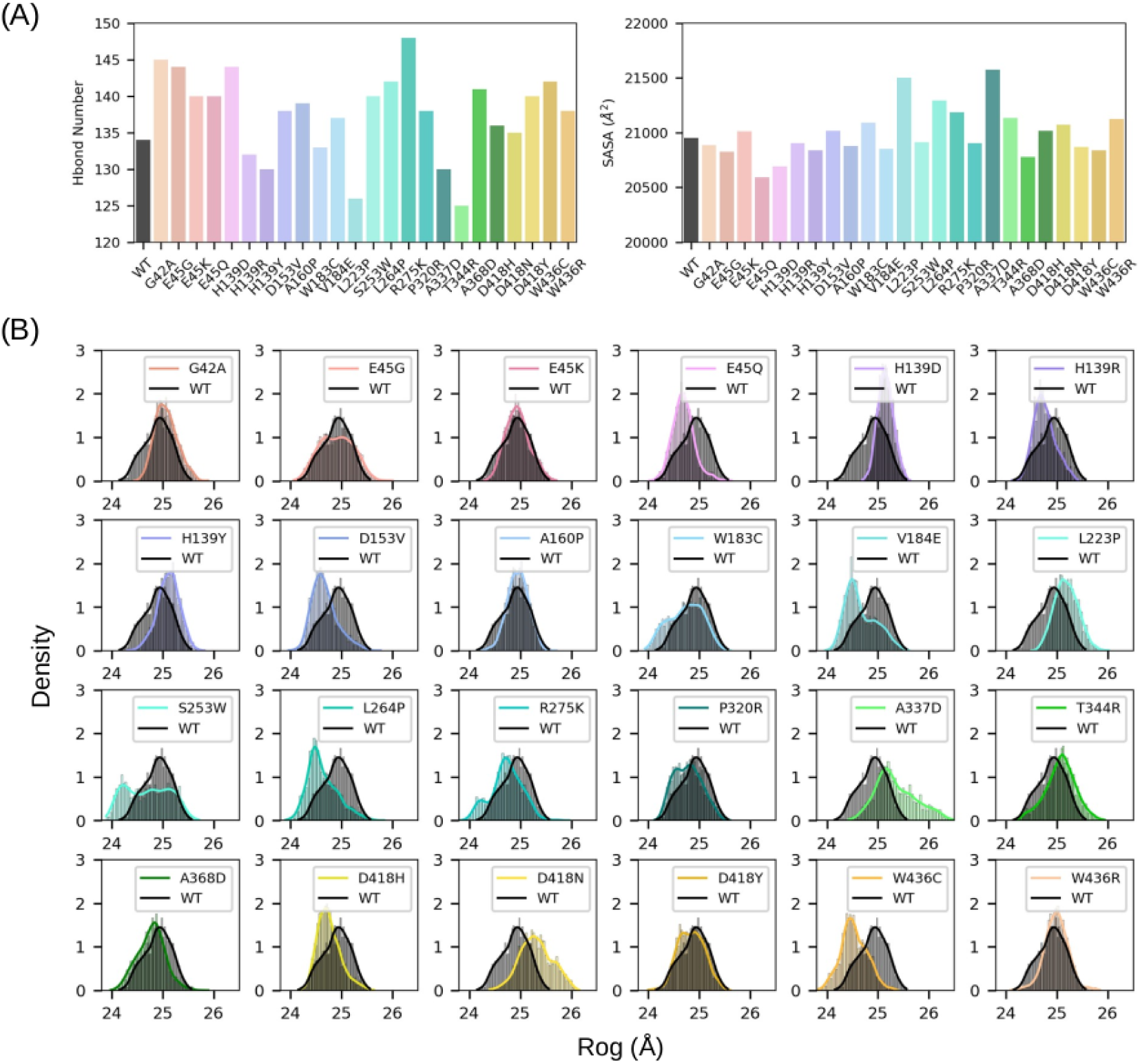
A)The average number of hydrogen bonds that have > 0.5 frequency sampled in the 500 ns MD. The average SASA (solvent accessible surface area) are slso shown. B) Distribution of radius of gyration (RoG) values of WT and mutants.

**Figure S2:**
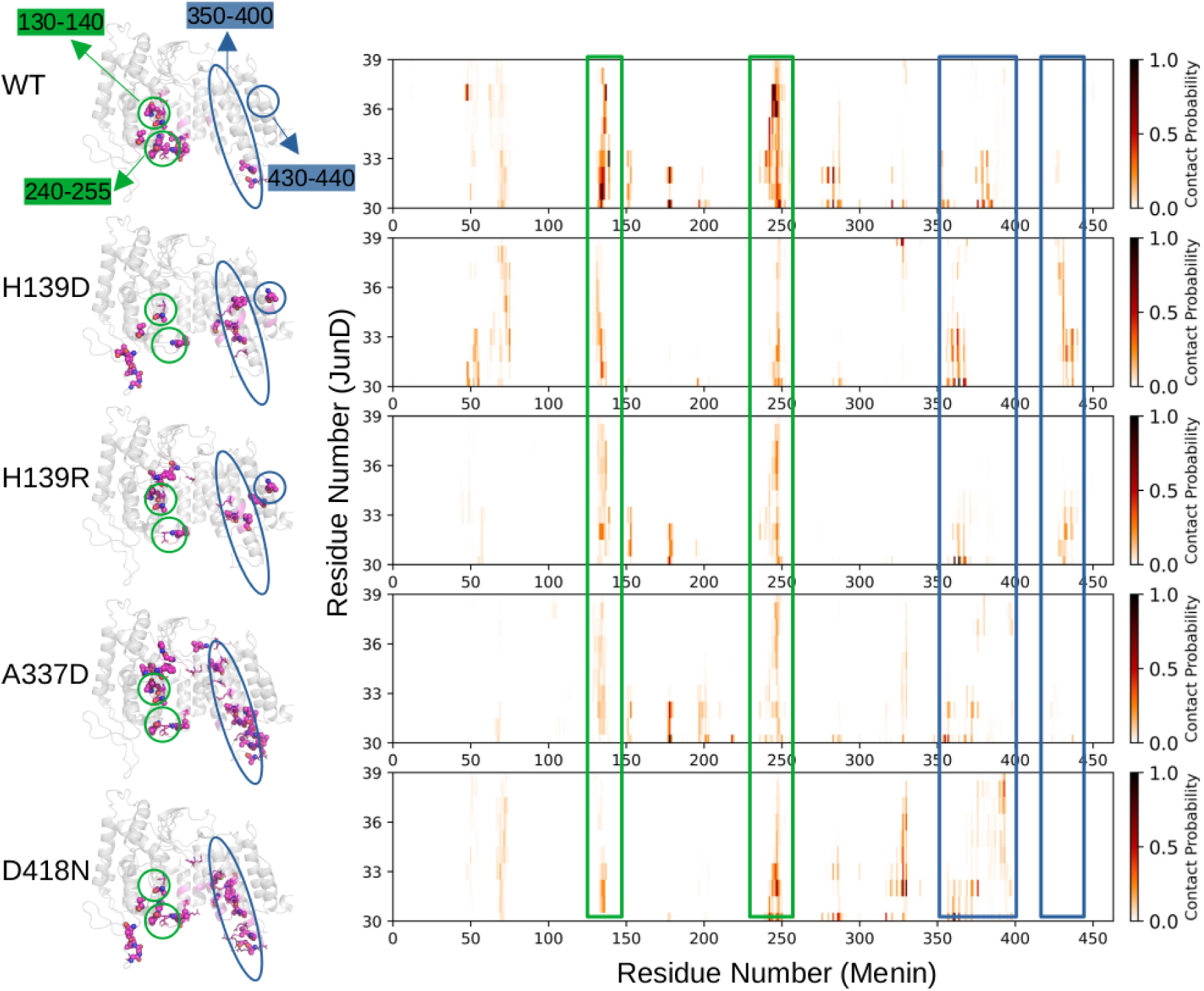
Mutations alter the regions where Menin interacts with JunD during the unbinding process. Menin residues that exhibit significant contacts with JunD during the umbrella sampling are colored magenta, with hydrophobic residues shown as spheres and alkaline residues shown as sticks. The contact probability heat map between JunD and Menin is shown on the right. Green boxes highlight regions where WT Menin exhibits higher contact frequency with JunD compared to the mutants, while blue boxes indicate regions where the mutants have higher contact frequency. These data demonstrate that in WT Menin, JunD primarily passes through regions comprising Menin residues 130-140 and 240-255 during unbinding. In contrast, in mutants H139D, H139R, A337D, and D418N, the contact frequency of JunD with these regions is diminished. Instead, the mutants interact with Menin in alternative regions: H139D and H139R primarily interact with JunD through regions 350-400 and 430-440, while A337D and D418N predominantly interact with JunD through region 350-400.

**Figure S3:**
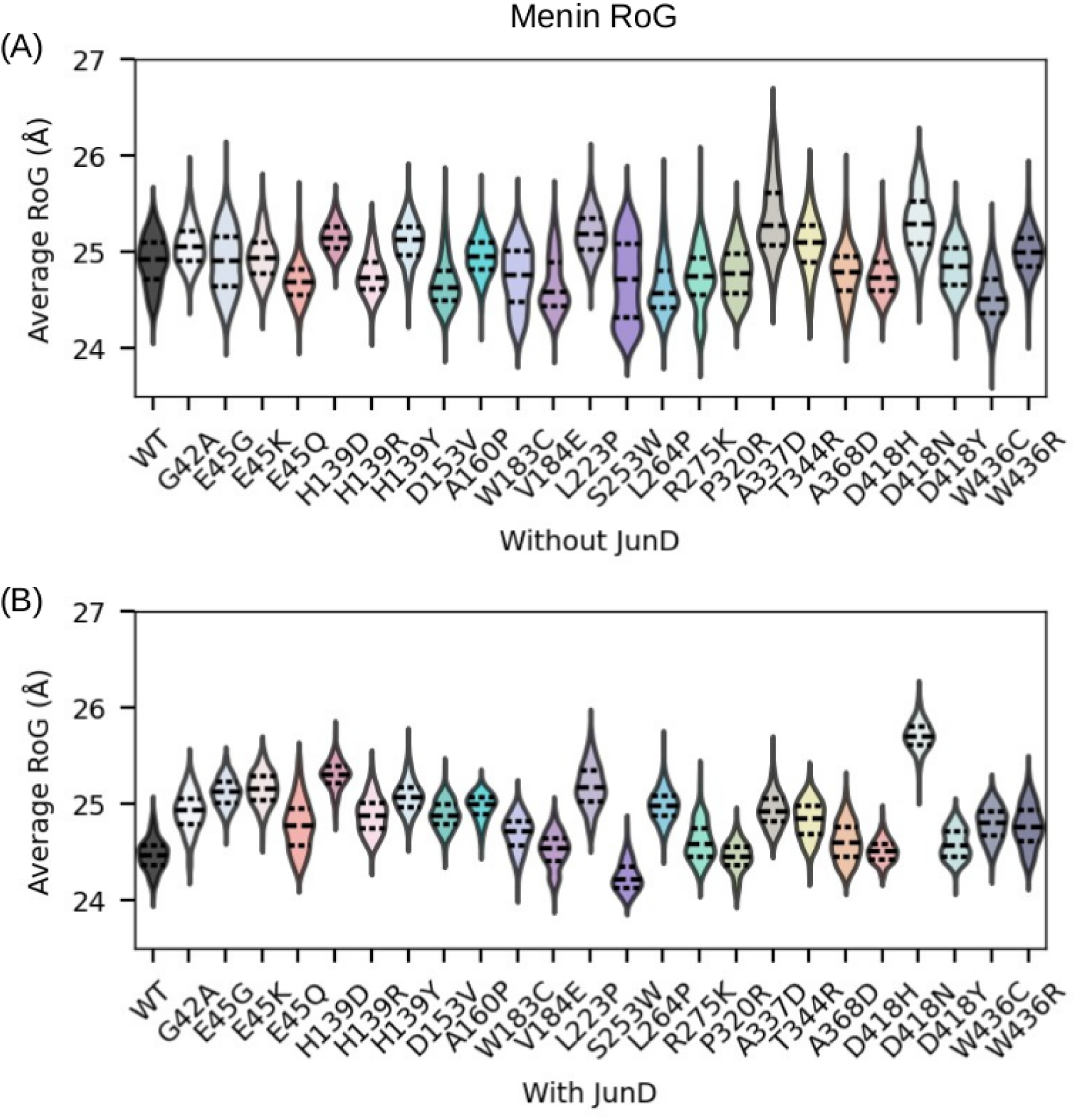
The presence of JunD affects the radius of gyration (RoG) distribution of Menin.

**Figure S4:**
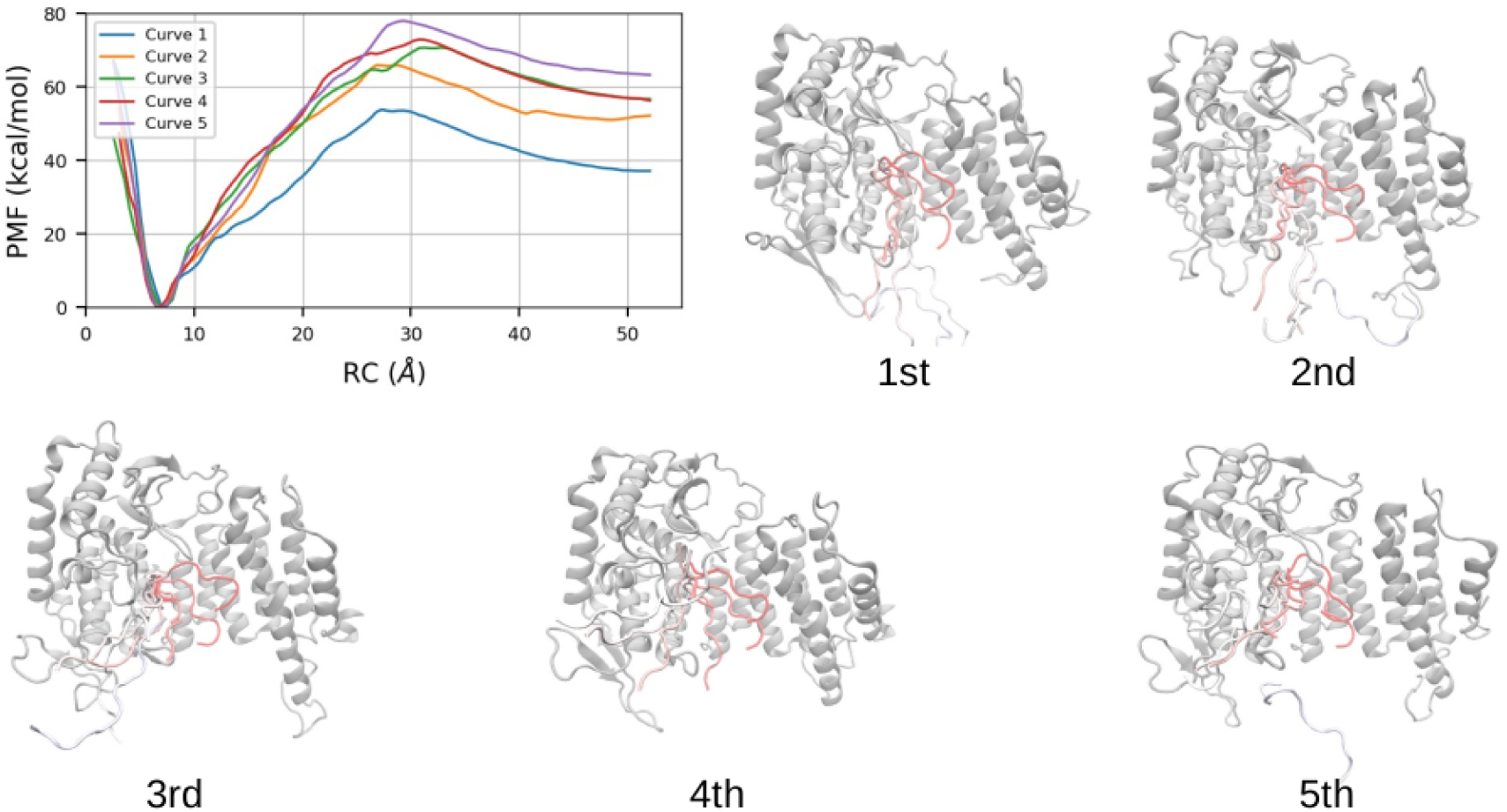
PMF curves of five independent replicas of umbrella sampling for the WT case. The dissociation pathway is depicted as window-dependent position of JunD with red color represents the initial window (bound state).

**Figure S5:**
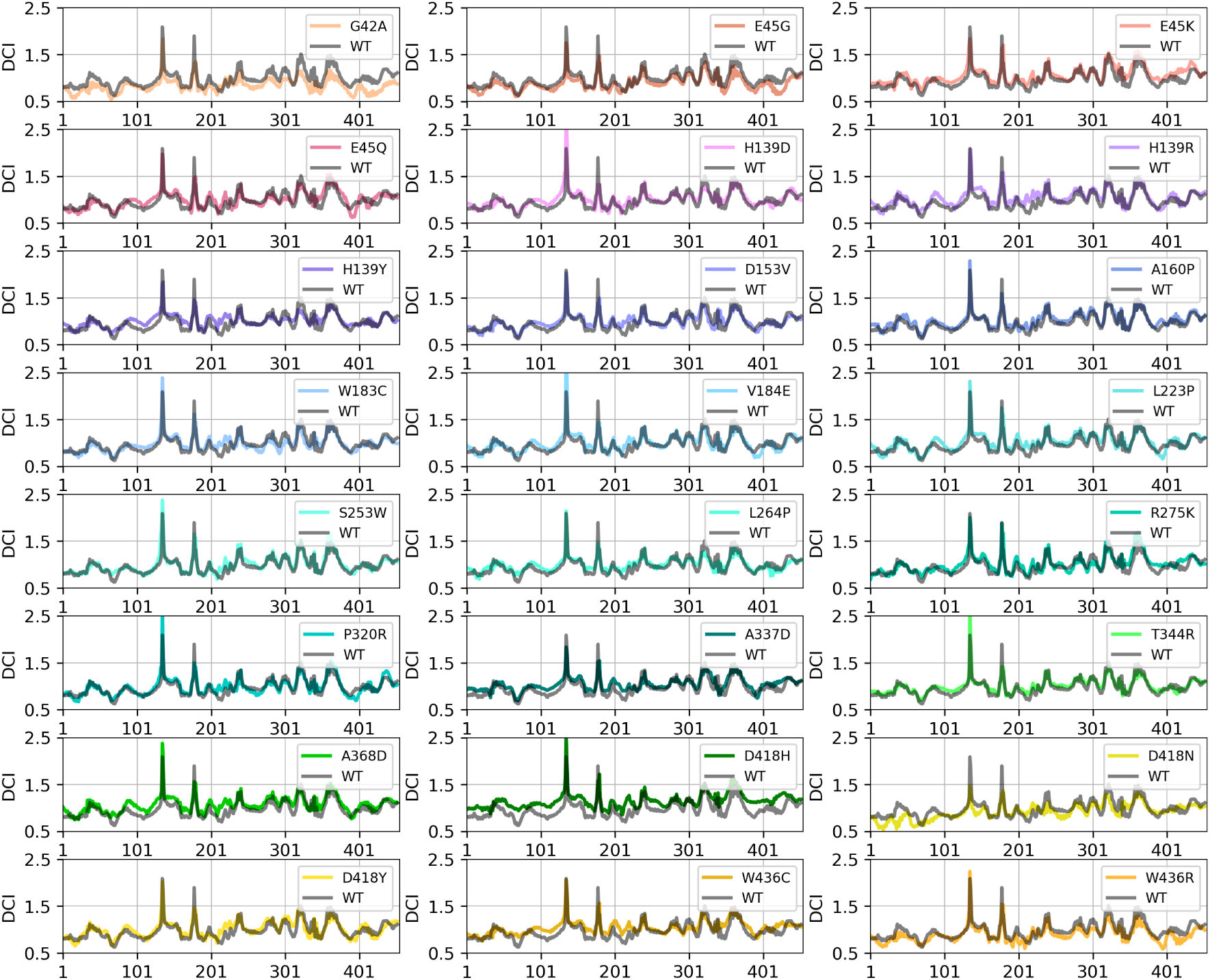
The per-residue DCI curve for each individual Menin mutants.

**Table S1:** All molecular dynamics simulations performed in the present study. Cases conventional MD Umbrella Sampling Rosetta-based conformation energy calculation: ROSETTA3="/opt/rosetta_src_2021.16.61629_bundle/main/source" # the structure to be scored is "input.pdb" $ROSETTA3/bin/relax.linuxgccrelease -s input.pdb -relax:ramp_constraints false\-relax:constrain_relax_to_start_coords -ex1 -ex2 -use_input_sc -flip_HNQ\-no_optH false -overwrite

| Cases | conventional MD |  | Umbrella Sampling |  |
| --- | --- | --- | --- | --- |
|  | Without JunD | With JunD | Without E179 restraint | With E179 restraint |
| WT | 500 ns | 100 ns | 5 ns/win×100 win×5 replica | 5 ns/win×100 win×5 replica |
| G42A | 500 ns | 100 ns |  |  |
| E45G | 500 ns | 100 ns |  |  |
| E45K | 500 ns | 100 ns |  |  |
| E45Q | 500 ns | 100 ns |  |  |
| H139D | 500 ns | 100 ns | 5 ns/win×100 win×5 replica | 5 ns/win×100 win×5 replica |
| H139R | 500 ns | 100 ns | 5 ns/win×100 win×5 replica | 5 ns/win×100 win×5 replica |
| H139Y | 500 ns | 100 ns |  |  |
| D153V | 500 ns | 100 ns |  |  |
| A160P | 500 ns | 100 ns |  |  |
| W183C | 500 ns | 100 ns |  |  |
| V184E | 500 ns | 100 ns |  |  |
| L223P | 500 ns | 100 ns |  |  |
| S253W | 500 ns | 100 ns |  |  |
| L264P | 500 ns | 100 ns |  |  |
| R275K | 500 ns | 100 ns |  |  |
| P320R | 500 ns | 100 ns |  |  |
| A337D | 500 ns | 100 ns | 5 ns/win×100 win×5 replica | 5 ns/win×100 win×5 replica |
| T344R | 500 ns | 100 ns |  |  |
| A368D | 500 ns | 100 ns |  |  |
| D418H | 500 ns | 100 ns |  |  |
| D418N | 500 ns | 100 ns | 5 ns/win×100 win×5 replica | 5 ns/win×100 win×5 replica |
| D418Y | 500 ns | 100 ns |  |  |
| W436C | 500 ns | 100 ns |  |  |
| W436R | 500 ns | 100 ns |  |  |

